# Proactive selective attention across competition contexts

**DOI:** 10.1101/2024.02.06.579112

**Authors:** Blanca Aguado-López, Ana F. Palenciano, José M.G. Peñalver, Paloma Díaz-Gutiérrez, David López-García, Chiara Avancini, Luis F. Ciria, María Ruz

**Affiliations:** Mind, Brain and Behavior Research Center (CIMCYC), University of Granada, Granada 18071, Spain; Department of Management, Faculty of Business and Economics, University of Antwerp, 2000, Belgium; Data Science & Computational Intelligence Institute, University of Granada, CP 18071, Spain

**Keywords:** biased competition model, selective attention, MVPA, preparation, EEG

## Abstract

Selective attention is a cognitive function that helps filter out unwanted information. Theories such as the biased competition model (Desimone & Duncan, 1995) explain how attentional templates bias processing towards targets in contexts where multiple stimuli compete for resources. However, it is unclear how the anticipation of different levels of competition influences the nature of attentional templates, in a proactive fashion. In this study, we used EEG to investigate how the anticipated demands of attentional selection (either high or low stimuli competition contexts) modulate target-specific preparatory brain activity and its relationship with task performance. To do so, participants performed a sex judgement task in a cue-target paradigm where, depending on the block, target and distractor stimuli appeared simultaneously (high competition) or sequentially (low competition). Multivariate Pattern Analysis (MVPA) showed that, in both competition contexts, there was a preactivation of the target category to select with a ramping-up profile at the end of the preparatory interval. However, cross-classification showed no generalization across competition conditions, suggesting different preparatory formats. Notably, time-frequency analyses showed differences between anticipated competition demands, reflecting higher theta band power for high than low competition, which mediated the impact of subsequent stimuli competition on behavioral performance. Overall, our results show that, whereas preactivation of the internal templates associated with the category to select are engaged in advance in both competition contexts, their underlying neural patterns differ. In addition, these codes could not be associated with theta power, suggesting different preparatory processes. The implications of these findings are crucial to increase our understanding of the nature of top-down processes across different contexts.

## 1. Introduction

In our everyday life we are surrounded by myriads of stimuli, but only some of them occupy our mind. The process of selective attention relates to the filtering of unwanted information and the selection of the pieces that are relevant to us. This intricate cognitive function proceeds through various biasing routes: one involves bottom-up processes, where the characteristics of stimuli automatically capture attention, while another is guided by goals, following top-down processes (Desimone & Duncan, 1995). The latter does not only take place during stimulus processing but can also happen in anticipatory fashion, by activating goal-related information before target events. This preparatory selection is related to proactive cognition (Braver, 2012). However, research about how such preparation unfolds to aid selection in contexts with different attentional demands is scarce.

One of the most influential proposals explaining selective attention is the biased competition model (Desimone & Duncan, 1995). This framework highlights the limited capacity of neural information processing and the essential role of competition in resolving this problem. As we move up along the cortical hierarchy, neurons increase their receptive fields to respond to stimuli. However, given the limits of their response, the more pieces of information (e.g., objects) are placed in the same receptive field, the less information there will be about each of them. Thus, neurons are selective and prioritize certain types of information, and thus the information that reaches our senses competes to be represented. Such competition is biased by bottom-up and top-down mechanisms. While bottom-up mechanisms may favor, for example, the most salient stimulus, top-down processes engage neurons in the prefrontal cortex that generate internal templates representing relevant information. These templates are used to bias neural competition favoring goal-relevant information. For example, if we aim to recognize a friend’s face in a crowd, pre-activation of a template of that face would later guide the attentional selection.

Early studies explored how top-down biases affect spatial selection (Moran & Desimone, 1985; Richmond et al., 1983) during target processing. To do this, authors compared the neural responses to stimuli in contexts of differential competition manipulated by the presence or absence of distractors. The first insights were obtained from electrophysiological cell recordings in non-human primates. For example, Moran and Desimone (1985) examined neurons of the visual cortex in a spatial attention task where monkeys had to respond to target stimuli placed at specific locations. They used stimuli with features that were effective or ineffective for a particular cell response. When these stimuli were presented simultaneously in the receptive field of the cell and the monkey attended to the effective stimulus, the neuron’s response was enhanced, whereas it was attenuated when the animal attended to the ineffective stimulus. Therefore, the cell’s response was determined by the properties of the attended stimulus. Later on, these findings were supported by data in humans using functional magnetic resonance imaging (fMRI). Kastner et al. (1998) studied how competition is resolved in the human brain when multiple stimuli appear simultaneously (a high competition context) or sequentially (low competition). They found that in conditions of inattention, a high competition context generated less activity in the visual cortex compared to a sequential presentation, indicating suppression of activation due to competition. Crucially, when the competition was biased by top-down spatial attention, neural suppression was reduced, corroborating the idea that focused attention magnifies attended information by mitigating the suppression caused by nearby stimuli. More recent studies have shown that selection also results in a heightened representation of specific characteristics of the attended stimuli (Kaiser et al., 2016; Reddy et al., 2009; Sheldon et al., 2021). These findings have been possible thanks to the use of advanced analytic techniques such as multivariate pattern analysis (MVPA), which has allowed to detect how brain activity patterns encode templates of attended features (Jackson et al., 2017) or categories (Kaiser et al., 2016; Reddy et al., 2009) of stimuli. Moreover, other studies have revealed changes in oscillatory activity, especially on the theta band. An increase in theta power has been found in midfrontal regions during target processing when conflictive stimuli compete with the target (Chevalier et al., 2021; Nigbur et al., 2011, 2012). Overall, this literature shows how stimulus selection in competition contexts is a complex process in which different neural mechanisms take part.

The studies discussed so far did not address neural processes that may be engaged during the preparation when the presence of upcoming competing stimuli can be anticipated. Some other studies have focused on preparatory activity (González-García et al., 2016; Peelen & Kastner, 2011; Peñalver et al., 2023; Rajan et al., 2021) but did not explore how varying levels of competition might influence specific preparatory processes. A fruitful approach to investigate preparation uses anticipatory cues to track preparatory templates, aligning with the principles of biased competition theory. This way, it has been found that both spatial and content-based information is preactivated before the target presentation (Rajan et al., 2021). For example, Peelen and Kastner (2011) used symbolic cues to instruct participants to detect either people or cars on naturalistic images entailing high levels of competition. Using MVPA, they compared the activity patterns in the visual cortex during the preparatory interval and during visual processing of exemplars from the target categories. Their results showed shared neural codes across both epochs, suggesting that an attentional template similar to the one guiding visual processing was preactivated during the preparation, biasing the competition in favor of stimuli matching this template. More studies have also found preparatory templates associated with relevant target categories (González-García et al., 2016; Peñalver et al., 2023; Ruz & Nobre, 2008) or stimulus features (Stokes et al., 2009) to attend. In a complementary manner, time-frequency analyses have revealed that theta power during the preparation period is associated with anticipating most challenging tasks (Cooper et al., 2017; Van Driel et al., 2015). Moreover, the behavioral relevance of both MVPA and time-frequency results has been evidenced by showing how both indices correlate with performance. On one hand, previous studies have shown a behavioral improvement when preparatory activity patterns associated with the target are more segregable (González-García et al., 2017; Peelen & Kastner, 2011; Soon et al., 2013; Stokes et al., 2009), when the dimensions to attend are better distinguished (Hall-McMaster et al., 2019) or when the working-memory load of the task is better represented (Manelis & Reder, 2015). On the other hand, preparatory theta power in frontocentral electrodes has been related to a more consistent behavior on task-switching paradigms (Cooper et al., 2017) or to a necessary step to accurate fast responses (Formica et al., 2022). However, the relationship between anticipated coding of specific information across contexts, theta power and behavioral performance remains uncertain.

In this work we examined if and how preparatory neural signals (i.e., the presence of target-specific activity patterns and theta band power increases) are affected by anticipated competition levels, as well as the relationship among them and with task performance. To do so, we collected EEG data during a cue-target paradigm with different levels of competition across blocks, including a separate localizer task to isolate perceptual templates. We analyzed anticipatory neural activity with univariate (time-frequency) and MVPA approaches. Considering previous findings (Hall-McMaster et al., 2019; Manelis & Reder, 2015; Peelen & Kastner, 2011; Peñalver et al., 2023), we expected that preparatory patterns would dissociate based on the relevant target category, and that this category-specific pattern would be more distinguishable in a high competition context. Also, at the oscillatory level we predicted that the amplitude of preparatory theta power would be enhanced in high competition (Cooper et al., 2017; Van Driel et al., 2015). Finally, we hypothesized that these preparatory brain signals would be related to behavioral performance (Formica et al., 2022; González-García et al., 2017; Peelen & Kastner, 2011; Soon et al., 2013; Stokes et al., 2009).

## 2. Methods

### 2.1. Participants

Thirty-six students (mean age = 21.36; range = 18-27; 18 women and 18 men) from the University of Granada, all native Spanish speakers, right-handed and with normal or corrected vision, were recruited and gave their informed consent to participate. They received 20-25 euros depending on their task performance. We excluded three additional participants due to either low accuracy (lower than 80%) or more than 30% discarded EEG trials due to artifacts. Data were collected during the COVID-19 pandemic; therefore, participants’ temperature was measured upon arrival, they wore a face mask during the experiment and signed a form confirming not having illness symptoms.

We calculated the sample size using PANGEA (Power ANalysis for GEneral ANOVA designs; Westfall, 2016). Our task followed a 3-factor within-subjects design (Competition x Stimulus Category x Congruency) where our main contrast of interest was a two-way interaction (Competition x Congruency). To achieve an estimated 80% power to detect a small-medium behavioral effect size of Cohen’s d = 0.3, we required a minimum of 30 participants. Nonetheless, to match counterbalancing needs we collected data from 36 participants, with an estimated power of 87%.

### 2.2. Apparatus, stimuli and procedure

The task was run on Matlab 2020a using The Psychophysics Toolbox 3 (Brainard, 1997). Stimuli were presented on an LCD screen (1920 × 1080 resolution, 60 Hz refresh rate) over a grey background. We used four types of stimuli as cues: circle, square, drop and diamond with thin black outlines, unfilled. As targets and distractor stimuli we employed 24 Caucasic faces with neutral expressions from the Chicago Face Dataset (Ma et al., 2015) and 24 Spanish person names (50% male-female in both categories).

When participants arrived to the lab they signed an informed consent, and then the EEG preparation started. They read the instructions of the task and performed a practice session (192 trials identical to the main task), where they had to achieve 80% of accuracy on both High and Low competition blocks to continue with the experimental session.

The experiment consisted of two tasks presented on different blocks: a main competition task and a stimulus category localizer. The main task was a cue-target paradigm where participants judged the sex of target faces and words. Cues presented at the beginning of each trial indicated the category of the target (faces/names) to respond to. Target and distractors were displayed either simultaneously (in High competition blocks, 50%) or sequentially, with a temporal delay (Low competition, 50%; adapted from Kastner and colleagues, 1998). Target and distractor stimuli could be either congruent (i.e., same sex, associated with the same response, 50%) or incongruent (different sex, with different responses, 50%). At the beginning of each block, an instruction screen stated the level of competition (High vs. Low) and indicated the cue-target associations for the block. To prevent perceptual confounds in the multivariate analyses, each category (faces and words) was cued with two different stimuli for each participant. That is, two cues always indicated faces, and the other two names. One of each pair was used in each block (one for faces and another one for names). Within participants, we counterbalanced the combination of cues across blocks, sequentially iterating across all possible pairs of face and name cues. The association between cues and target categories was further counterbalanced across participants.

The sequence of events in a trial was as follows (see Figure 1): The cue (∼2°x 2° degrees of visual angle) was presented for 50 ms and was followed by a Cue-Target-Interval (CTI) of 1500 ms. In High competition blocks, an overlapping face (∼9.7° x 12.17° visual angle) and a name (∼9.7° x 2.6° visual angle) used as target and distractor stimuli were displayed for 750 ms, followed by an Inter-Trial-Interval (ITI) of 1500 ms. In Low competition blocks, the target appeared first on the screen for 500 ms, followed by overlapping target and distractor for 250 ms and then by the distractor on its own for 500 ms, ending with a 1000 ITI ms. This arrangement follows previous similar paradigms (see Kastner et al., 1998) and allows to present each stimulus with the same duration and maintain the same trial length across competition conditions. The response window was the same in both conditions. Participants pressed the keys “A” or “L” with their left and right index to indicate whether the target stimulus was female or male (counterbalanced across participants). In case of wrong answers, after the ITI, a feedback tone of 450 Hz was played for 300 ms while a fixation cross was displayed for 1000 ms in total.

**Fig. 1.**
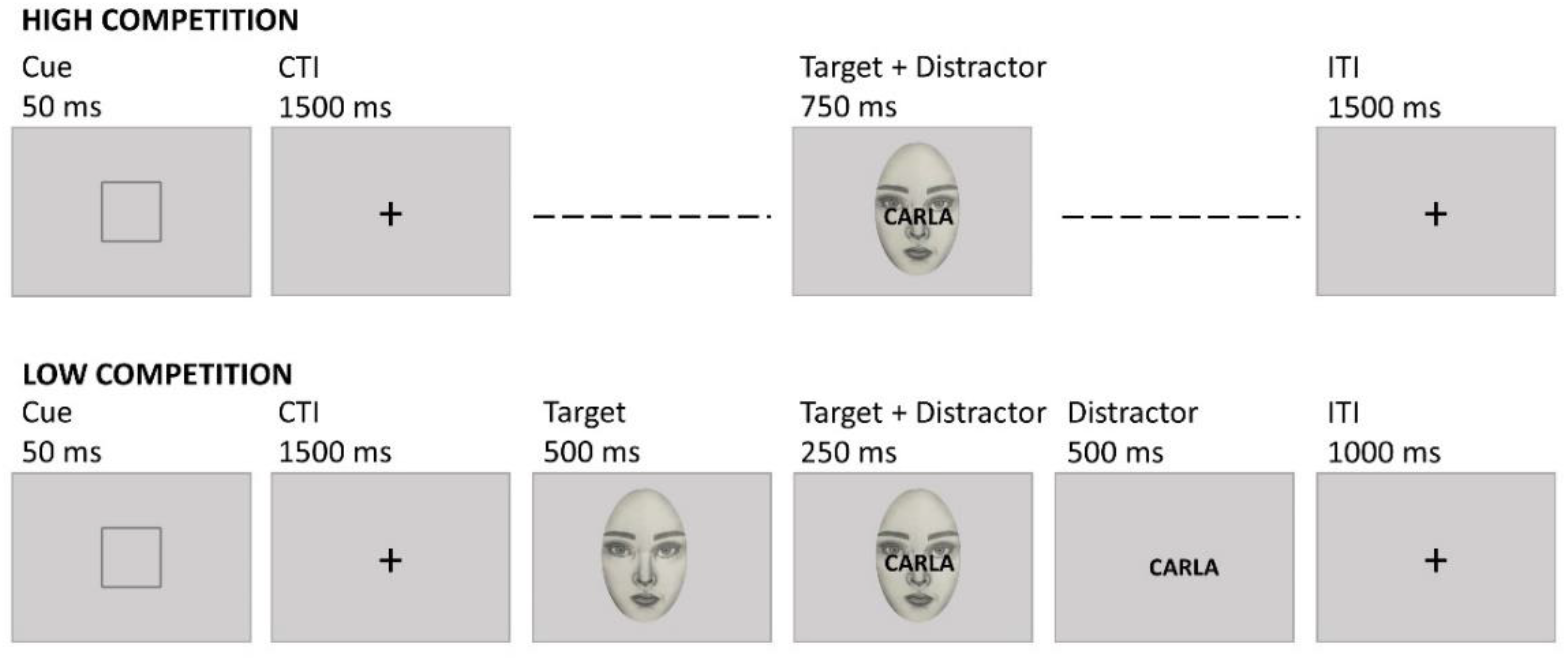
Experimental paradigm. Example trials from High and Low competition blocks. In a sex classification task, participants were cued about which stimulus category (faces or names) to respond to. In High competition blocks, targets and distractors appeared at the same time, whereas in Low competition they appeared sequentially.

**Fig. 2.**
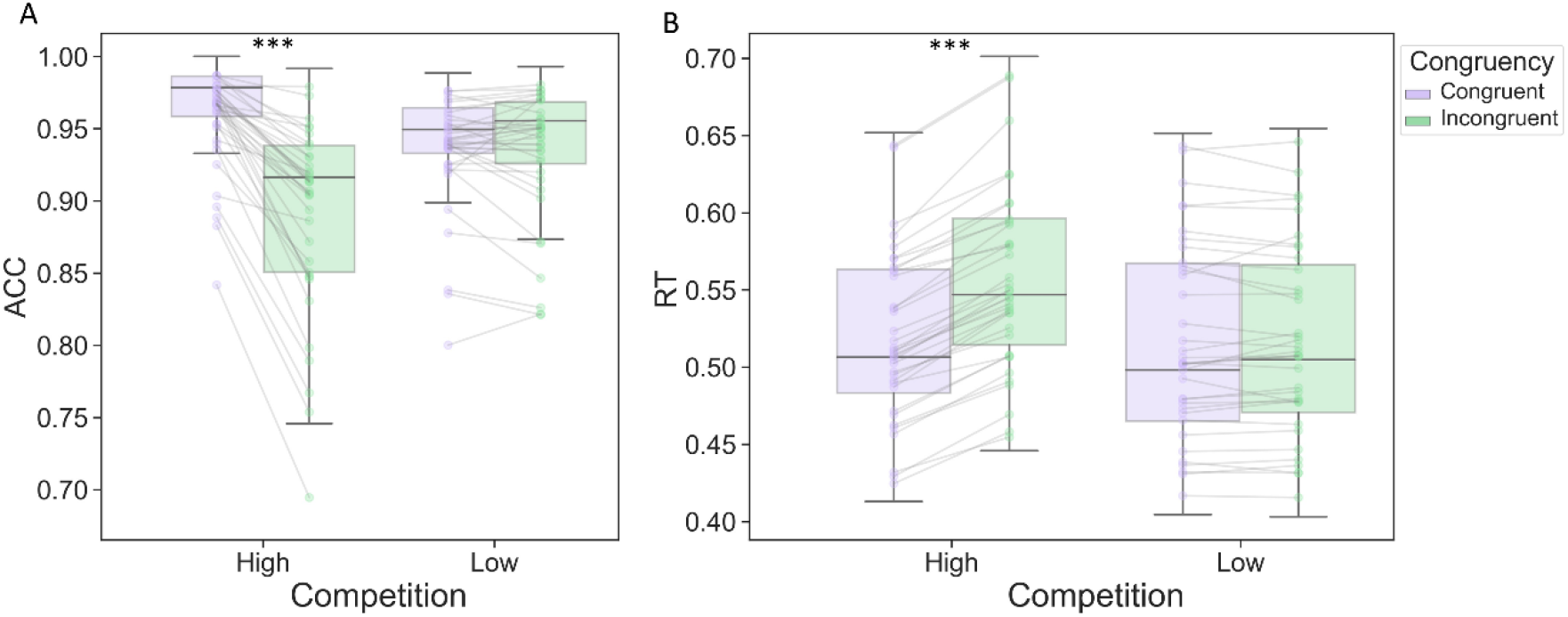
Box plots displaying the behavioral results. Boxes have a middle marking the median, limits representing the first and third quartile, and whiskers indicating the 1.5 inter quartile range for the upper and lower quartiles. Outliers are shown outside the whiskers. The dots represent each participant’s value per experimental condition. (A) Behavioral accuracy rate (ACC) in High and Low competition blocks for Congruent and Incongruent trials. (B) Reaction times (RT in seconds) in High and Low competition blocks for congruent and incongruent trials. *** = *p* < 0.001

In Localizer blocks, included to isolate the perceptual processing of stimuli without motor activity, the same faces and names were presented for 750 ms followed by an ITI of 500 ms. To facilitate participants’ engagement with the task, they were instructed to press the "C" key in a minimal percentage of trials (8%) where the target was rotated 180°.

There were 72 blocks, 24 of each type (High competition, Low competition and Localizer). The order of blocks was fully counterbalanced within and across participants, as each block was preceded and followed by the other types the same number of times. For the main task, we had 576 trials for each (High, Low) competition condition. These trials were unique combinations of the 24 faces and 24 names. Each block lasted 1.52 minutes (with 24 trials of 3.8 s each). The localizer blocks had 48 trials lasting 1.25 s, for a total of 1 minute per block (for a total of 1152 localizer trials). The whole session, including practice, lasted approximately 2 hours and 15 minutes.

### 2.3. EEG acquisition and preprocessing

EEG data was recorded with a high-density 64 active channels cap (actiCap Slim, BrainVision) at the Mind, Brain and Behavior Research Center (CIMCYC) of the University of Granada. The impedances of the amplifier were kept below 10 k Ω. EEG activity was recorded at a sampling rate of 1000 Hz, with FCz as the reference electrode.

Preprocessing was done using EEGLAB (Delorme & Makeig, 2004) and in-house MATLAB scripts following the pipeline available on Github (see Open practices section). First, data were downsampled to 256 Hz and filtered using a low-pass and high-pass FIR at 120 and 0.1 Hz, respectively. A notch filter was applied at 50 and 100 Hz to remove line noise and its harmonics. Noisy channels were identified by visual inspection and removed (1 channel on average, range 0-4). Next, the data was epoched in intervals of 3 s (-1 to 2 s after the onset of cues and of targets). Independent Component Analysis (ICA) was computed afterwards with the runica algorithm from EEGLAB to remove blinks and lateral eye movements. Components were selected with ICLabel and visual inspection (scalp maps, raw activity and power spectrum). An average of 1.58 components per participant (range 1-3) were removed. Then, automatic trial rejection was used to prune the data from other artifacts, using 3 criteria (see López-García et al. (2020, 2022) and Peñalver et al. (2023) for similar parameters). First, we identified trials with abnormal spectra, removing those deviating from baseline by ± 50 dB in the 0–2 Hz frequency window (sensitive to remaining eye artifacts) or by -100 dB or + 25 dB in 20–40 Hz (sensitive to muscle activity). Second, trials with improbable data were eliminated: the probability of occurrence of each trial was computed by determining the probability distribution of voltage values across trials, with a rejection threshold established at ± 6 SD. The third criteria were extreme values: all trials with amplitudes in any electrode out of a ± 150 μV range were rejected. Next, the dismissed channels were recomputed by spherical interpolation and a common average was used to re-reference the data. Finally, we applied a baseline correction in the -200 to 0 ms prior to stimulus onset. The analyses focused solely on correct trials. On average, 1476 trials per participant (range 1292-1591) were included.

### 2.4. Analyses

#### 2.4.1. Behavioral

We employed 2-way repeated measures ANOVAs with the factors Competition (High vs. Low) and Congruency (Congruent vs. Incongruent). Separate tests were performed on accuracy and reaction times (RT) using the JASP software (Love et al., 2019). To filter the RT data, we excluded incorrect trials and those with RT deviating 2SD from the participant mean.

#### 2.4.2. EEG

##### 2.4.2.1. Multivariate pattern analysis (MVPA)

We used Linear Discriminant Analyses (LDA) as classifiers to investigate if the preparatory activity patterns contained information about the upcoming competition level (High or Low) and the specific target categories (Faces or Names) anticipated across competition contexts. To do so, we focused on the cue-locked interval activity from -100 to 1550 ms. The analyses were run on MATLAB using the toolbox MVPAlab (López-García et al., 2022). Classifiers were trained and tested using raw voltage of each trial and time point across all the channels, with the configuration for the classification being equal for all the analyses.

To increase the signal-to-noise ratio, we created ‘supertrials’ (Grootswagers et al., 2017) by averaging three random trials within each condition (see López-García et al. (2020, 2022); Peñalver et al. (2023) for similar procedures) and smoothed the data by applying a moving average window every three time points, so that data from every timebin (t_n_) was averaged with the previous and the following time-points t_n_ = (t_n-1_ + t_n_ + t_n+1_)/3. We used a 5-fold cross-validation strategy that ensures unbiased results while reducing the computational cost (Grootswagers et al., 2017). With this approach, we split our data into five subsets and used four to train the classifier and the remaining one as a test set. This protocol was repeated 5 times, changing the test set. The number of trials within each class was subsampled considering two criteria: that each class had the same number of trials and that there was the same number of trials per class in each fold for the cross-validation procedure (Grootswagers et al., 2017; King & Dehaene, 2014). A normalization procedure was applied to enhance the classifier performance and generalizability of the results. Normalization was done during the cross-validation, by calculating the mean and standard deviation of each electrode within each fold across the training trials, and then applying these two values to normalize the data of both the train and the test set as:

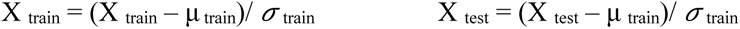

Where μ _train_ is the mean and *σ* _train_ is the standard deviation of the training set. Finally, to reduce the computational cost, the analysis was done every three time points. The results are reported as the area under the curve (AUC), a non-parametric criterion-free method with no assumptions about the true distribution of the data (King & Dehaene, 2014). Also, AUC is sensitive to binary (two-class) differences, less vulnerable to biases (including those triggered by potential differences between the classes) and can be interpreted as classification accuracy (King et al., 2013).

To detect the significant decoding performance at the group level, we used a non-parametric cluster-based permutation method against empirical chance. For this, the trial labels were randomly permuted 100 times per participant, resulting in chance-level outcomes under the null effect. After that, one random AUC per participant was selected and averaged to create a group-level null effect decoding curve. This was done 10^5^ times to generate 10^5^ permuted group AUC values. These values were used to build empirical chance-level AUC distributions for each time point. The AUC values in the 95 percentiles were used as threshold to identify significant decoding peaks in the real decoding results. Moreover, to estimate the minimum cluster size to be significant (α = 0.05), we used the permuted results to generate a null distribution of cluster sizes and corrected for multiple comparisons using a False Discovery Rate (FDR) approach (López-García et al., 2022).

Additionally, we studied the extent to which the activity patterns were stable along the preparation interval. For that purpose, we used a temporal generalization approach that applied the decoding analysis explained above but training in a given time point and testing in all the remaining ones. This procedure iterated using all time points as training and testing datasets. This resulted in a Temporal Generalization Matrix with a diagonal reflecting the same result as the MVPA curve and non-diagonal values corresponding to the temporal generalization of the underlying neural code. The statistical significance from these matrices was extracted with the same cluster-based analysis as before, now considering two-dimensional clusters that spread over training and testing time points.

With this overall approach, we first studied if the competition level affected preparatory activity. To do so, we trained and tested classifiers on the cue-locked interval of trials from High and Low competition blocks. Then, to evaluate whether and how the preparatory interval carried information about the target category anticipated, either faces or names, we performed classification analyses separately for High and Low competition blocks. To compare the category-related patterns while avoiding perceptual confounds triggered by the specific shape of the cues, we adopted a cross-classification approach (Kaplan et al., 2015; Peñalver et al., 2023). The classifiers decoding the relevant category were trained and tested in trials where independent sets of cues were employed. This protocol iterated across the two sides of the classification (exchanging training and testing cues) and all cues’ combinations (e.g.: circle-names, diamonds-faces vs. drops-names, squares-faces). Results were averaged across directions and classifiers.

Additionally, we further studied if category-specific coding was affected by the competition level anticipated in High and Low competition contexts, following two approaches. First, we tested the hypothesis that preparing for high competition contexts increased the fidelity of anticipatory category-specific neural codes. To do this, we compared the two competitions cross-decoding curves and temporal generalization matrices using a tailored cluster-based permutation approach (Moore et al., 2024). We computed one-tailed *t*-tests in every time point to address whether the decoding accuracy or the temporal generalization was higher in High than Low competition contexts. Then, we identified cluster sizes in these results, by looking for sets of temporally adjacent points with *p* < 0.05. For that purpose, we set a criterion of a minimum cluster size of 3 time points the cross-decoding curve, and 10 for temporal generalization (Peñalver, 2024). To each of these clusters, we assigned a *t* value resulting from the sum of the *t* values from all the time points incorporated. Next, the inference was performed contrasting these results against a permutation-based distribution of null differences. To do so, we randomly multiplied one of the conditions by -1 in each subject; this was done 5000 times. We repeated the procedure as in the true data but with the permuted results, obtaining a distribution of the *t* values of the clusters, and used the 95^th^ percentile to mark the values that were considered significant in the true data. Second, to further explore whether the competition context altered the neural codes underlying the anticipatory category patterns, we performed a cross-classification training and testing the classifier with data from different competition blocks (Kaplan et al., 2015). This analysis also implemented the cross-classification across cues, following a similar approach as above. The AUC curves obtained were averaged among classifiers and directions.

To study whether the preparatory activity patterns associated with categorical information of faces and names were similar to the actual perception of the stimuli, we performed a cross-classification analysis by training with data from the localizer blocks and testing on the cue-locked window of the main task, separately for High and Low competition. As the timing of the localizer and main task paradigm were different, we focused on the temporal generalization profile of the cross-classification. Considering that our interest was the reinstatement of perceptual patterns on preparation activity, this cross-classification was only performed in a single direction, using the localizer data as training set and the main task data as test set. To test whether the reinstatement has different robustness in High or Low-competition contexts, we compared these matrices using two-tailed *t*-tests with a cluster-based permutations approach. This analysis was equivalent to the one comparing the category-specific coding across competition conditions, except for using two-tailed tests.

Finally, to study the link between anticipatory activity patterns and task performance, decoding results and behavioral data were correlated using the Pearson coefficient. First, we used the anticipated competition level, extracting the average AUC value for each participant during the time window where the decoding was significant at the group level, from 100 ms until the end of the interval (see Li et al., 2022). This value was correlated with behavioral accuracy and RT means across High and Low competition. To check for specific relationships between the congruency behavioral effect and the anticipated competition level, we calculated differences in task performance (separately for behavioral accuracy and RT) of congruent minus incongruent trials and correlated this with individual AUC values. Second, we followed a similar strategy with the fidelity of the category-specific decoding, but in this case, the AUC values of each participant were calculated separately for High and Low competition blocks. We averaged the AUC values during the time window that was significant in both High and Low competition contexts (1150-1550 ms). Then, we correlated the AUC with the behavioral accuracy and RT of each competition condition. To address the relationship with the congruency effect, we calculated the difference between the congruent and incongruent trials of each competition condition separately and correlated them with the mean AUC values of each condition. In all cases, we applied frequentist and Bayesian statistics to provide complementary evidence supporting the results.

##### 2.4.2.2. Time-frequency analysis

We tested whether preparing for High competition increased anticipatory theta-band activity (3-7 Hz) in comparison with Low competition. Theta power was extracted from a frontocentral region of interest (ROI) with the electrodes Fz, FC1, Cz, FC2, F1, C1, C2, F2 and FCz. We computed the time-frequency decomposition for each trial during the preparation epoch (cue locked -1000 to 1550 ms) using complex Morlet wavelets. The frequencies were logarithmically spaced in 18 steps from 2 to 20 Hz. The wavelet’s length was calculated separately for each frequency assigning a number of cycles also logarithmically spaced between 3 and 5 (see Cohen, 2019). Time–frequency power values were transformed to decibels and normalized to a baseline of -280 to -100 ms before cue onset, according to the following equation (Cohen & Van Gaal, 2014): dB= 10* log10 (power/baseline).

A cluster-based nonparametric statistical test implemented in FieldTrip (Maris & Oostenveld, 2007) was used to evaluate whether the preparatory activity of High competition trials showed higher theta power than Low competition ones. For this, power values in each condition were averaged across channels and trials. Then, these averages were compared using within-subjects paired-samples two-tailed *t*-test for each time point and frequency (Hz). Those *t* values larger than the threshold specified by alpha (0.05) were clustered in connected sets of temporal adjacency. The *t* value of the cluster was calculated adding the *t* values of each timepoint. The permutations were performed within each subject randomizing the condition labels for each value, 1000 times with the Monte Carlo method. *T* values were calculated for all the permutations using maximum cluster-level mass statistic (Groppe et al., 2011), and the most extreme cluster-level *t* score across permutations was used to derive a null hypothesis distribution. If the *t* value of the true data cluster was above the 97.5^th^ percentile or below the 2.5^th^ percentile of the null distribution, then it was considered significant.

Next, to explore whether anticipatory theta power increase and the content-specific activity patterns found with the decoding were related, we correlated them using Pearson across participants, separately for High and Low competition conditions. To obtain the theta power values per participant, we visually inspected the grand average across trials from all conditions and identified the theta time window from 100 ms to 900 ms (see Fig S1). We extracted the average theta amplitude per participant from this time window. Then, these values were correlated with the classification AUC per participant and competition condition averaged within the 1150 ms to 1550 ms time window.

Finally, we performed a mediation analysis to investigate if the anticipatory theta power acted as a mediator between the competition manipulation and the speed of responses (e.g., Formica et al., 2022). To do so, we used trial-by-trial RT and theta power data, averaging the power values within 100-900 ms after cue presentation (same time window as above). To filter out outlier data, trials with ± 2SD from the average RT or theta power were discarded. Afterwards, we verified that the data fitted the necessary criteria (Baron & Kenny, 1986) for mediation analysis: in our case (1) the competition manipulation had to influence RTs, (2) this manipulation also had to predict theta power, and (3) theta power had to predict RTs. These were tested with linear mixed effects models (LMMs) with the lme4 package in R (Bates et al., 2014). In all models, we included Congruency as a fixed effect to control for it. To select the model with an adequate random effect structure, a “keep it maximal” approach was adopted starting with the most complex random structure until the model converges for the 3 LMMs (Barr et al., 2013). This approach gave a random structure of (competition | subject). P-values were calculated using Satterthwaite approximations (Luke, 2017). Once we ensured the three criteria were met, the mediation was tested performing a casual mediation analysis with the function mediate from the mediation package in R, using 5000 permutations (Tingley et al., 2014).

## 3. Results

### 3.1. Behavioral

Overall accuracy on the main task was 93.5%. The ANOVA of accuracy data showed a main effect of Competition (F_35,1_ = 16.022, *p* < 0.001, η_*p*_^2^ = 0.314), indicating that participants performed better on Low (M = 93.8%, SD = 4.4%) than High competition blocks (M = 92.3%, SD = 5.2%). The Congruency effect (F_35,1_ = 71.269, *p* < 0.001, η_*p*_^2^ = 0.671) was also significant, showing that Congruent trials were more accurate (M = 95.1%, SD = 3.8%) than Incongruent ones (M = 91.2%, SD = 5.8%). As predicted, there was a significant interaction of Congruency * Competition (F_35,1_ = 73.030, *p* < 0.001, η_*p*_^2^ = 0.676). Post-hoc test showed that the Congruency effect appeared in High (M _Congr =_ 95.9%, M _Incongr_ = 88.6%, t_35,1_ = 9.319, *p* < 0.001, Cohen’s d = 1.553) but not in Low competition (M _Congr_ = 94.1%, M _Incongr_ = 93.6%, t_35,1_ = 1.555, *p* = 0.129, Cohen’s d = 0.259).

An average of 5.5% trials per participant, corresponding to outliers’ values of RT (± 2SD), were excluded. The ANOVA results showed a main effect of Competition (F_35,1_ = 26.664, *p* < 0.001, η_*p*_^2^ = 0.432), as participants were slower on High (M = 535.0 ms, SD = 60.0) compared to Low competition trials (M = 514.0 ms, SD = 68 ms). There was also an effect of Congruency (F_35,1_ = 224.819, *p* < 0.001, η_*p*_^2^ = 0.865) with faster Congruent (M = 515.0 ms, SD = 62.0) than Incongruent (M = 534.0 ms, SD = 63.0) trials, and an interaction of Congruency * Competition (F_35,1_ = 203.907, *p* < 0.001, η_*p*_^2^ = 0.853). Again, Congruency was only present in High competition (M _Congr_ = 517.0 ms, M _Incongr_ = 556.0 ms, t_35,1_ = -19.021, *p* < 0.001, Cohen’s d = - 3.170) and not in Low competition blocks (M _Congr_ = 514.0 ms, M _Incongr_ = 514.0 ms, t_35,1_ = 0.065, *p* = 0.949, Cohen’s d = 0.011).

### 3.2. Electrophysiology

#### 3.2.1. MVPA results

##### 3.2.1.1. Anticipation of the competition level

Our first aim was to assess the anticipation of the overall competition level. A classifier trained and tested to discriminate preparatory activity between High and Low competition contexts showed an effect of competition. The cluster identified in these results covered most of the CTI (from 100 ms until the end of the interval, see Fig. 3A). The temporal generalization analysis revealed a large significant cluster starting approximately at 100 ms. Interestingly, the cluster was asymmetric, generalizing to all the testing time points when the classifier was trained using data from the end of the preparation window, whereas less generalization was found when the training was done at the beginning of the interval (see Fig. 3B).

**Fig. 3.**
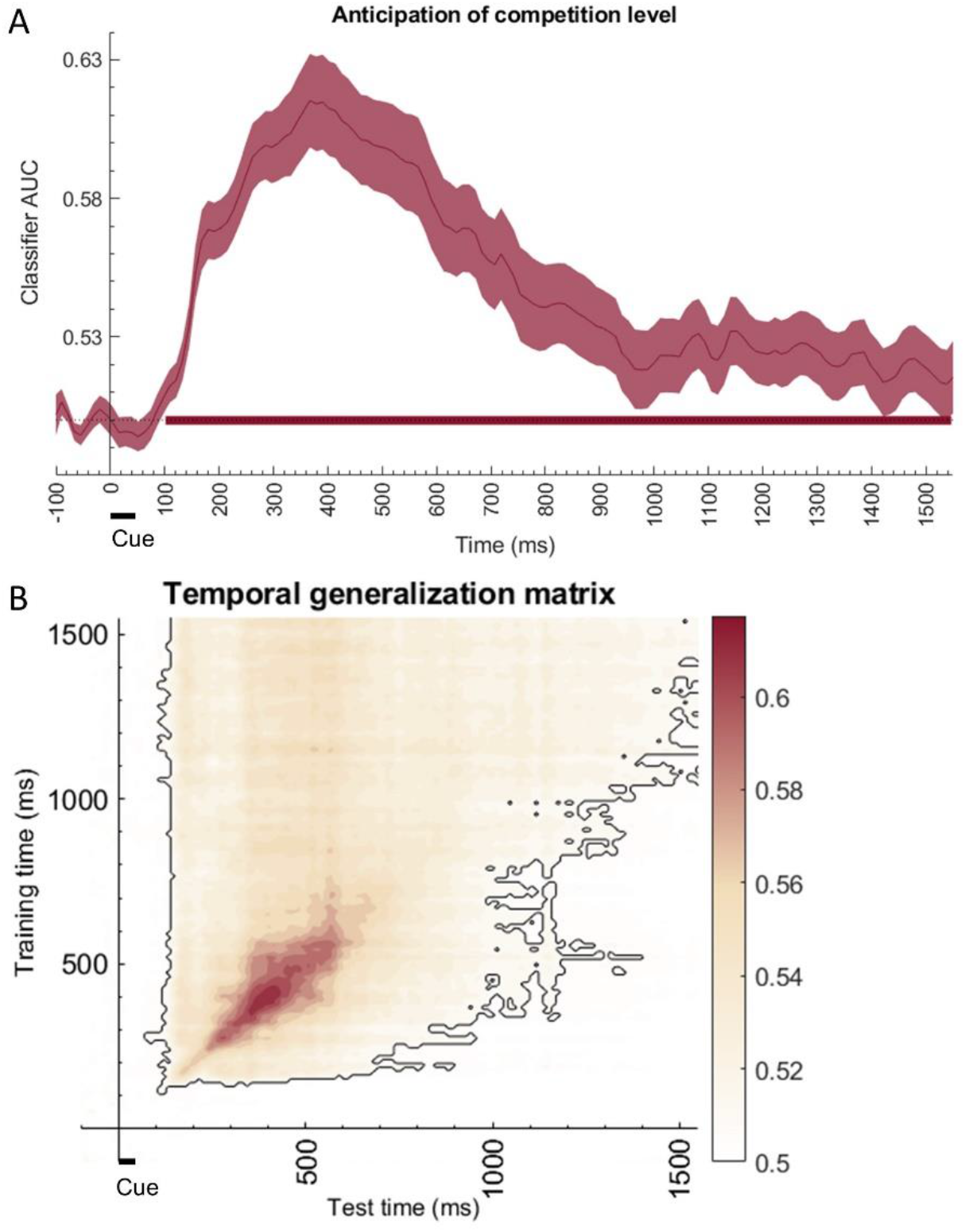
(A) Classifier performance distinguishing competition levels in the preparation interval. The red line shows the mean AUC and the shaded red areas its standard error. The red horizontal line displays significant time points. The black horizontal line shows the onset and duration of the cue. (B) Temporal generalization matrix from the same classification showing significant above-chance clusters outlined in black. The color range in the bar indicates the AUC values.

##### 3.2.1.2. Category-specific anticipation

Classifiers trained and tested to discriminate the target category that the participants were preparing to attend (faces vs. names) indicated that there was a significant effect of category. We found decoding clusters in High (from 531-555 ms, 859-906 ms, 976-1000 ms, 1047-1082 ms, 1152-1211 ms, 1234-1527 ms) and Low competition contexts (941-965 ms; 1176-1234 ms; 1281-1328 ms; 1352-1398 ms; 1527-1550 ms). The decoding AUC incremented progressively with a ramping up profile towards the end of the preparation interval, before target onset (see Fig 4). Nonetheless, there was no evidence supporting different decoding accuracies across competition conditions (all *ps* > 0.05).

**Fig. 4.**
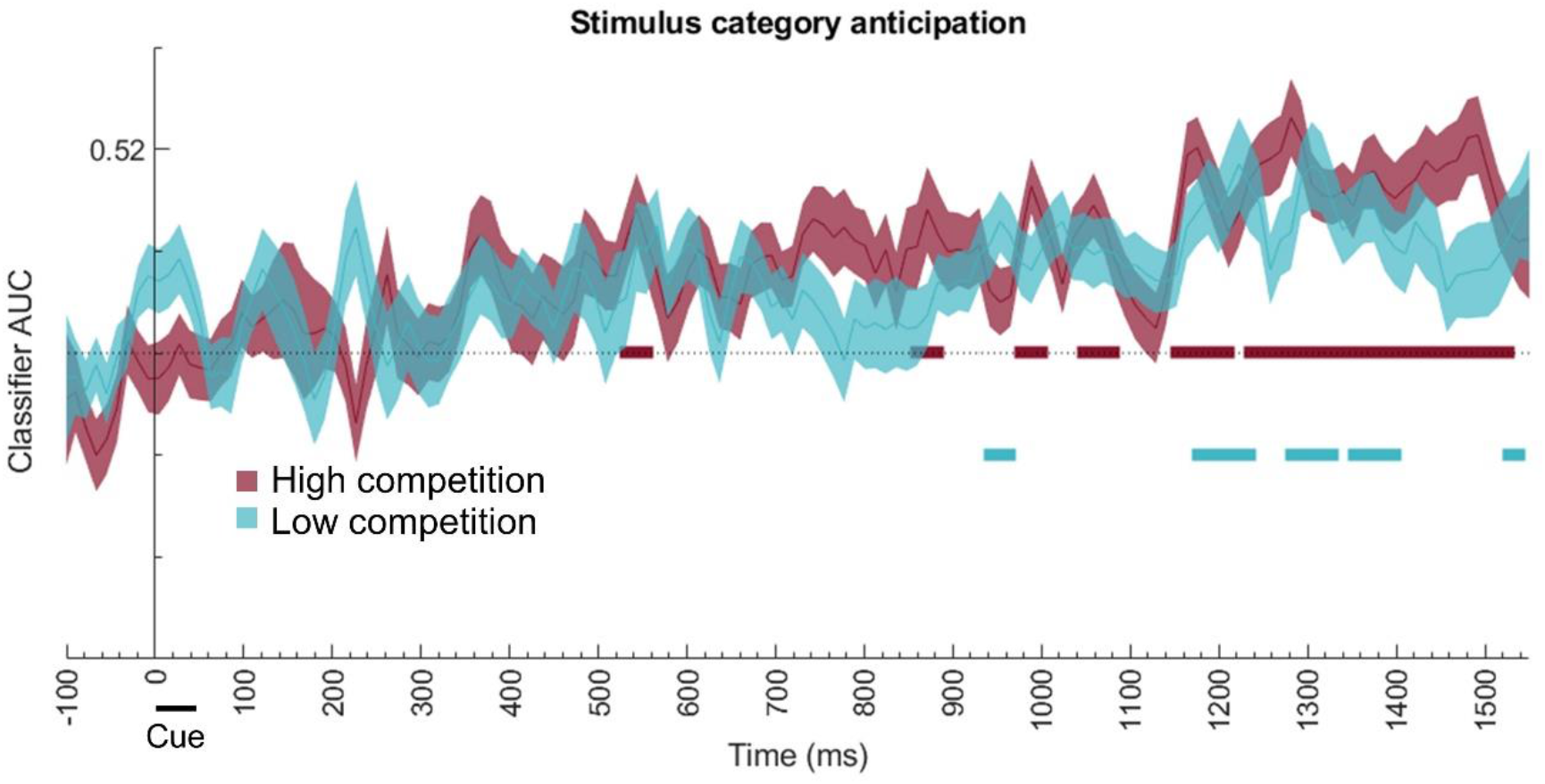
Results (AUC values) of the classifiers discriminating the upcoming target category (faces vs. names) using cross-classification (across cues identities), separately for High and Low competition conditions. Horizontal lines represent the significant clusters for High (red) and Low (blue) competition.

We also analyzed the temporal generalization of category-specific information separately for High and Low competition contexts. For both, the anticipatory patterns showed temporal generalization, which was stronger on the right upper corner of the matrix, towards the end of the interval (Fig. 5). However, the comparison of both matrices did not provide statistical evidence supporting different generalization patterns (all *ps* > 0.05).

**Fig. 5.**
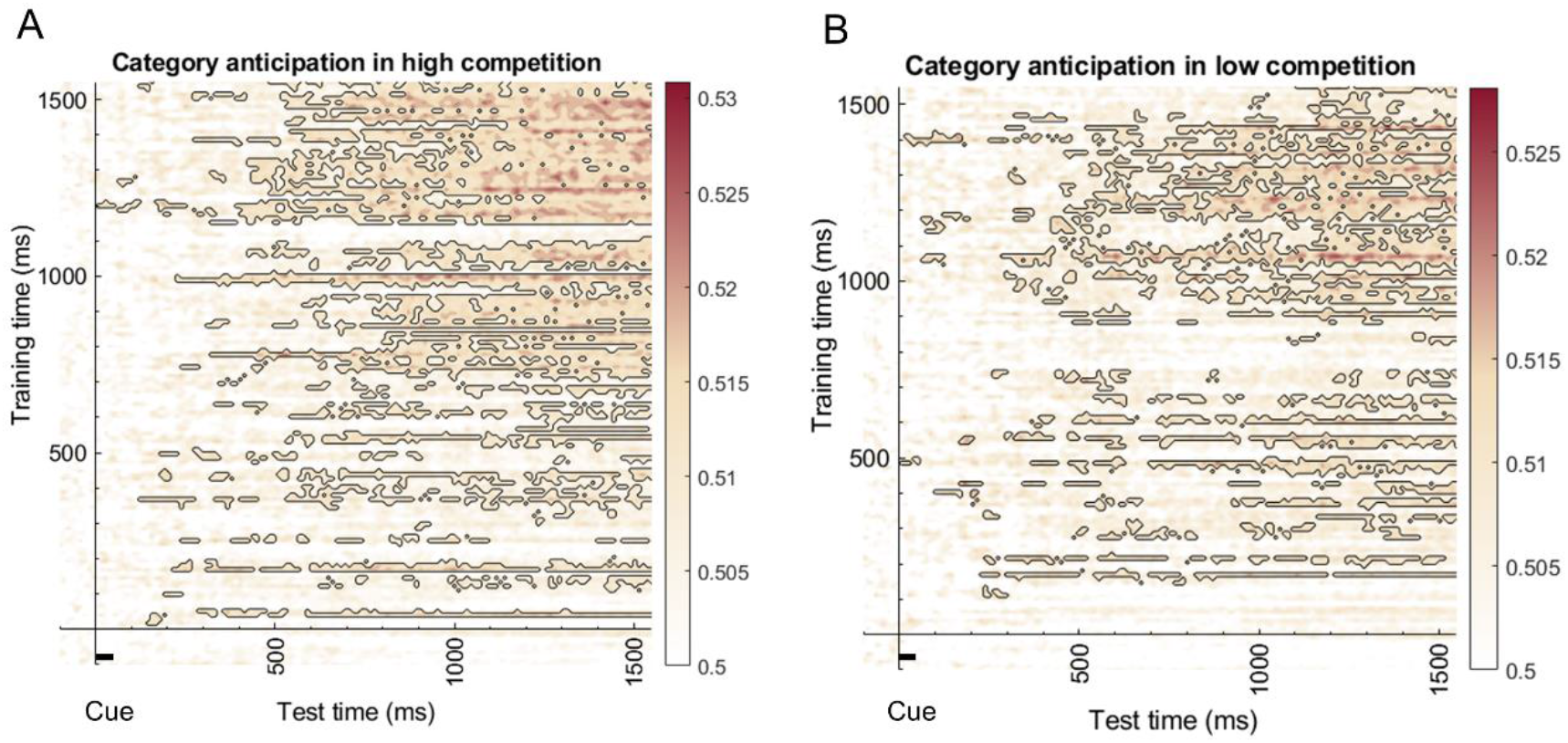
Time-generalization matrices of the discrimination of the upcoming target category (faces vs. names) using cross-classification (across cues identities) on High (A) and Low competition (B). Significant clusters above chance are outlined with black. The color range in the bar represents AUC values.

We also tested whether these patterns in High and Low competition are coded similarly. A cross-classification strategy across conditions and cues showed no evidence for similar patterns coding the relevant category between competition contexts (Fig. 6A). The temporal generalization analysis only showed small scattered significant clusters (see Fig. 6B).

**Fig. 6.**
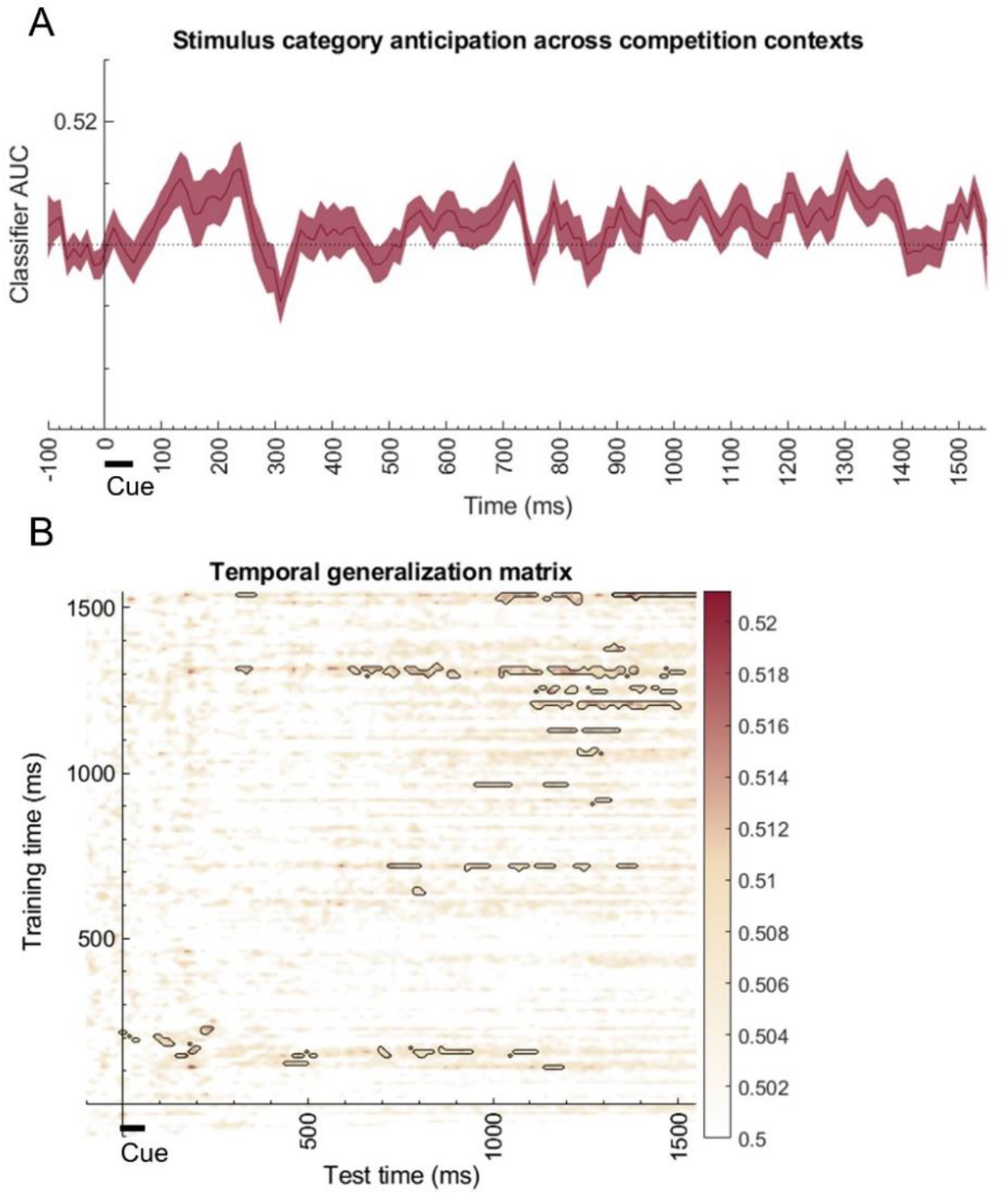
(A) Cross-classification across competition conditions and cues differentiating upcoming target categories, showing no significant clusters. (B) Temporal generalization matrix for the same cross-classification showing small clusters above chance, outlined with black. The color range represents AUC values.

##### 3.2.1.3. Preactivation of perceptual patterns during preparation

Classifiers were trained to discriminate between faces and names in the localizer and tested during the preparation interval of the main tasks separately for High and Low competition contexts. The results showed several above-chance significant clusters in the High competition context (Fig. 7A). In the preparation interval of Low competition trials, there were few above-chance clusters (Fig. 7B). However, when comparing both matrices, there was no statistical evidence supporting different preactivations of perceptual patterns across competition conditions (all *ps* > 0.05).

**Fig. 7.**
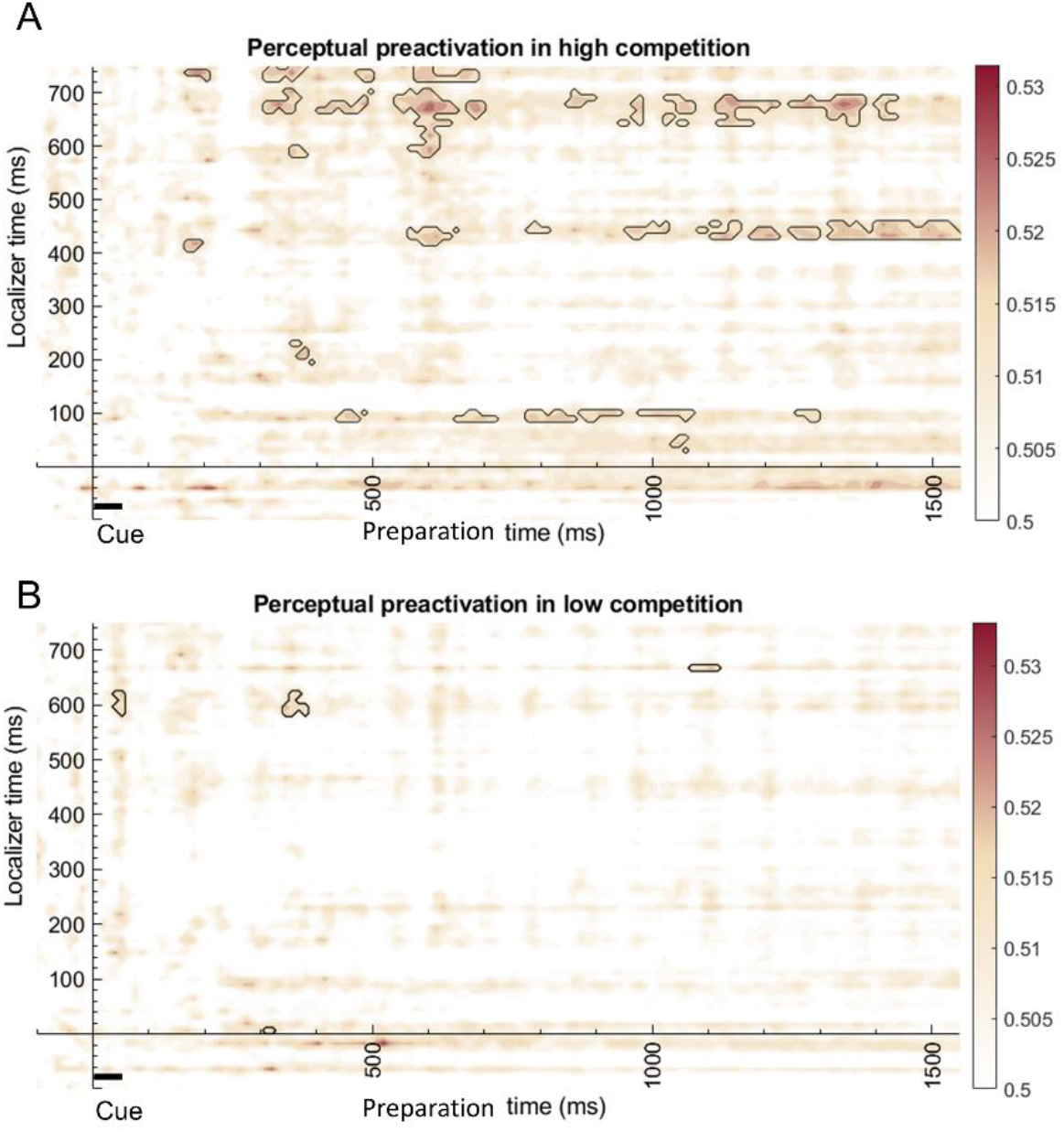
Temporal generalization of the cross-classification from the localizer category targets to the upcoming category targets in the preparation interval of (A) High competition and (B) Low competition (with cue duration presented as the black horizontal line). The color range in the bar represents AUC values.

##### 3.2.1.4. Decoding-behavior relationships

To study whether the observed preparatory patterns of competition were related to behavioral performance, we correlated the decoding accuracy of the classifier with the average accuracy and RT across participants. However, Pearson correlations resulted in non-significant results (all *ps* > 0.6). A Bayesian approach provided moderate evidence in favor of the null hypothesis, i.e., the absence of a relationship between the two variables (Mean behavioral accuracy: *r* = 0.03, *p* = 0.87, BF_01_ = 4.75; Mean RT: *r* = 0.09, *p* = 0.60, BF_01_ = 4.22). We also correlated the behavioral congruency effect (i.e., the accuracy and RT difference of congruent minus incongruent trials) and the anticipation of competition level decoding. Pearson correlations were non-significant (all *ps* > 0.4) and Bayesian factors showed moderated evidence towards the null hypothesis (Behavioral accuracy: *r* = -0.03, *p* = 0.83, BF_01_ = 4.71; RT: *r* = 0.11, *p* = 0.52, BF_01_ = 3.94).

We also correlated behavioral performance with category-specific decoding values separately for High and Low competition indexes. In this case, for each participant, we averaged the AUC in the same time window for High and Low competition. Again, the Pearson correlations were non-significant (all *ps* > 0.2) and Bayesian factors showed weak to moderate evidence in favor of the null hypothesis (High competition accuracy: *r* = 0.16, *p* = 0.34, BF_01_ = 3.12; High competition RT: *r* = -0.22, *p* = 0.20, BF_01_ = 2.15; Low competition accuracy: *r* = -0.01, *p* = 0.96, BF_01_ = 4.81; Low competition RT: *r* = -0.01, *p* = 0.97, BF_01_ = 4.82). A similar approach was taken to correlate the congruency effect with the category-specific anticipation decoding. The results indicated that none of the correlations were significant (all *ps* > 0.2), providing weak to moderate evidence in favor of the null hypothesis (High competition, accuracy: *r* = -0.03, *p* = 0.88, BF_01_ = 4.76; High competition, RT: *r* = 0.19, *p* = 0.27, BF_01_ = 2.68; Low competition, accuracy: *r* = 0.03, *p* = 0.86, BF_01_ = 4.75; Low competition, RT: *r* < 0.01, *p* = 0.98, BF_01_ = 4.82).

#### 3.2.2. Time-frequency results

The comparison of the two conditions’ time-frequency maps in the cue-locked interval showed that High competition anticipation generated higher Theta power than Low competition. A large cluster (*p* < 0.001, Fig. 8) was found around the Theta band (3-7 Hz) from 0 to 1550 ms.

**Fig. 8.**
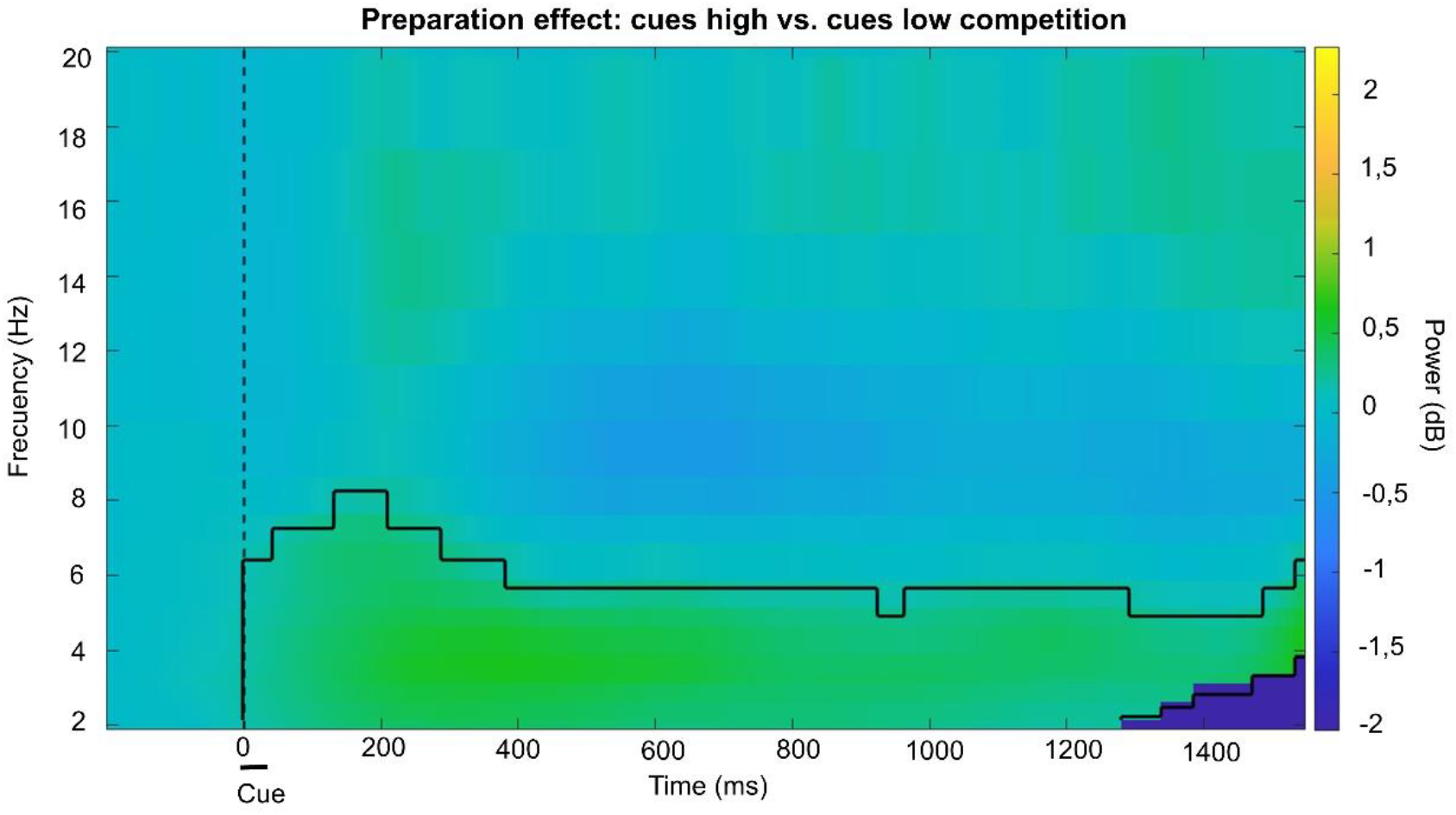
Results of the Monte Carlo cluster-based approach comparing the power values of High and Low competition trials during the preparation interval. The significant cluster is outlined with black lines. The dark blue patch on the lower-right corner reflects the lack of estimated power data due to edge effects.

The Pearson correlations between category-specific decoding and theta power in High and Low anticipation conditions resulted in non-significant results (all *ps* > 0.6). A Bayesian approach provided moderate evidence supporting the null hypothesis (High competition: *r* = 0.08, *p* = 0.65, BF_01_ = 4.37; Low competition: *r* = 0.08, *p* = 0.65, BF_01_ = 4.38).

Finally, we explored whether the neural mechanisms reflected by the frontocentral theta power could act as mediators between the impact of competition levels and the RTs. The filtering performed prior to the analysis removed an average of 7.5% (SD= 1%) of trials for each participant. We confirmed that our data met the necessary criteria for the mediation (Baron & Kenny, 1986), fitting our data on three LMMs (see Supplementary materials for details). First, in agreement with previous analysis, there was a Competition and Congruency effect on RTs. Second, the effect of Competition on theta was also significant, so that High competition induced higher theta power. Third, in the complete model predicting RTs there was an effect of Congruency, Competition and also theta on RTs, suggesting that larger theta values were associated with faster responses. To directly test that theta power partially mediates the effect of competition on RTs, we performed a causal mediation analysis that showed a significant direct effect of competition on RTs (β = 0.022, CI 95% = [0.013, 0.03], *p* < 0.001) and an indirect effect via theta (β = -0.0001, CI 95% = [-0.0002, 0], *p* = 0.006), indicating a partial mediation (see Fig. 9).

**Fig. 9.**
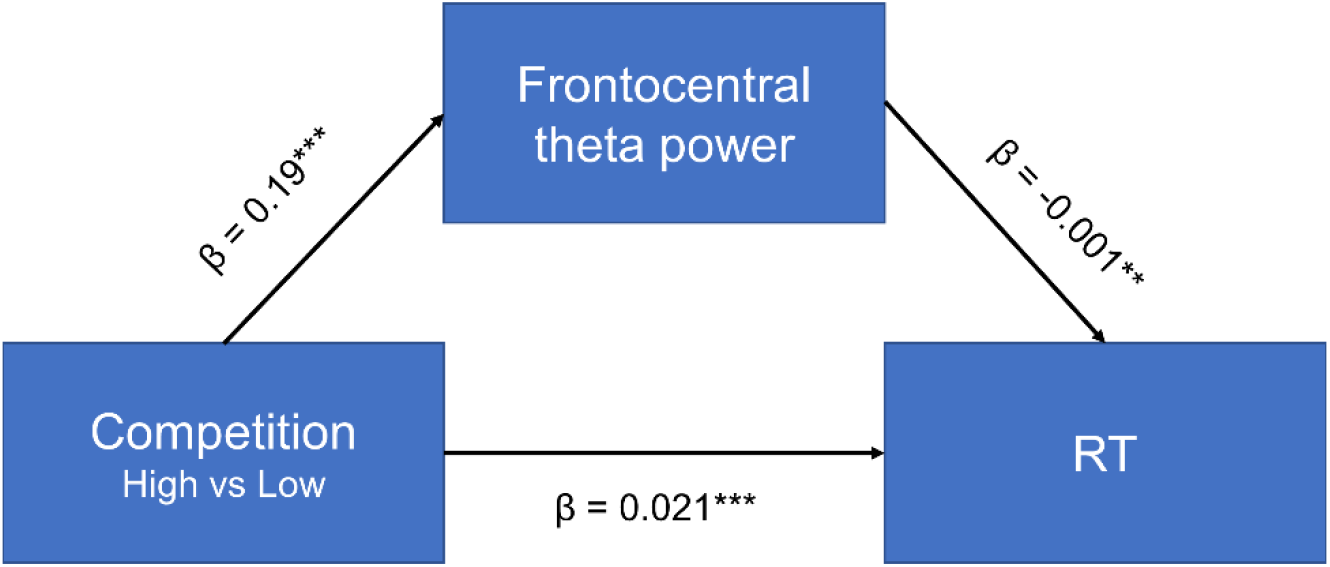
Mediation model with beta values. The anticipated competition significantly predicted frontocentral theta power, which in turn affected the RTs. Frontocentral theta power mediated the effect of competition on RT, although the direct effect of competition on RT was still significant after accounting for the mediating effect of theta, suggesting a partial mediation. \*\*\**p* < 0.001 \*\**p* < 0.01

## 4. Discussion

In this study we examined how the anticipated demands of attentional selection, manipulated through competition between target and distractors, modulated specific markers of preparatory neural activity. In line with the postulates of the biased competition model, our results reveal the preactivation of internal templates associated with the category to select. However, these preactivations did not differ in robustness across competition contexts, and they did not generalize between competition contexts, suggesting the existence of differential preparation formats. Moreover, oscillatory activity showed higher theta band activity for high than low competition context, an effect that mediates behavioral improvements.

Our behavioral results validated the effectiveness of the paradigm. The high accuracy rates across conditions show that participants paid attention to the cues, and the congruency effect was present only in the high competition condition, in line with classic studies (Beck & Kastner, 2009; Desimone & Duncan, 1995; Eriksen & Eriksen, 1974; Simon, 1969). The higher selective attention demands were also reflected in lower accuracy and higher response times in the high competition blocks, as expected (Desimone & Duncan, 1995; Duncan, 1993).

Multivariate classifiers showed that anticipated competition levels could be distinguished. These differences between high and low competition contexts are in line with previous studies that found evidence of preparatory coding of different tasks (González-García et al., 2017; Hall-McMaster et al., 2019; Manelis & Reder, 2015; Palenciano et al., 2019a). Importantly, the competition level could be differentiated in most of the preparation window. However, it is worth noting that these results could also be driven by other variables that may be systematically modulated by competition levels. That is the case for instance of arousal, which could be increased in high competition blocks. Also, our blocked design makes it highly likely that the control settings (i.e., the overall task set of high vs. low competition contexts) were maintained throughout the whole duration of the block (Dosenbach et al., 2008; Palenciano et al., 2019b). Further studies will be needed to disentangle the differences between control settings with various competition levels and arousal changes.

Unexpectedly, the temporal generalization profile of the overall competition context was asymmetric, showing more generalization when the classifier was trained at the end of the interval than for the reverse direction (Fig. 3B). This could be due to more stable preparatory patterns at the end than at the beginning of the interval. However, earlier time points still resulted in higher accuracy values, which may be caused by having a set of processes occurring at the same time, which might be different for high and low competition contexts. Some of these early cognitive processes might include physically perceiving the cue, remembering its meaning, recalling the competition condition in which the participant is and preparing to perceive the target and respond to it. As this set of processes might be taking place at the same time only at the beginning of the interval, generalization to the rest of the time window may not be possible. However, as the preparatory interval was fixed, participants could predict target appearance, therefore at the end of the preparation interval neural activity could reflect to a greater extent preparatory processes, such as category-specific patterns shown at the decoding associated with the target category. These patterns, observed upon completion of the interval, may also appear (alongside others) at different points within the preparation interval, thus exhibiting greater generalizability.

Our results also indicate the presence of specific preparatory patterns linked to the anticipated category of the target to select, both in high and low competition contexts. This is consistent with the theory of biased competition, as a reflection of an internal attentional template associated to the relevant category. Preparation is specific to the content of the incoming target to select (González-García et al., 2017; Palenciano et al., 2019a; Peelen & Kastner, 2011; Peñalver et al., 2023; Rajan et al., 2021; Sobrado et al., 2022; Stokes et al., 2009), with a strength of category anticipation increasing at the end of the interval, in a ramping-up fashion, replicating Peñalver and colleagues (2023). Although understanding the implications of this finding requires further research, it could be related to temporal expectations. As mentioned earlier, the preparatory interval was fixed, consequently the predictable temporal structure of the task could intensify the preactivation of specific stimulus patterns towards target appearance (Jin et al., 2020; Rohenkohl et al., 2012). Another possible, non-exclusive mechanism is the processing of the cue meaning. This may occur silently (Stokes et al., 2009) at the beginning of the preparation interval as evidenced in reduced decoding accuracy, and it may reactivate when needed just before the target. The temporal generalization matrices of the categories on both competition contexts also display a progressively increasing generalization toward the end of the preparatory interval, suggesting that the activity patterns were not only stronger, but also more stable over time at later stages (King & Dehaene, 2014).

The comparison of the preparatory category coding in high and low competition did not detect differences between conditions in either the fidelity of these patterns or their temporal stability. Hence, these results suggest that the categorical attentional templates were equivalent across competition contexts. Although unexpected, this result resonates with previous evidence showing that attentional templates also arise in low competition contexts (González-García et al., 2016). Given that predictive cues were used in both contexts, it is reasonable that the representation of target category was found in both situations. While the classifier accuracy remained undistinguishable across contexts, the anticipated competition introduces other differentiated process that might not be captured by the classifier alone, implying a multi-faceted approach to information representation. Critically, finding equivalent classification accuracies does not imply that the underlying neural codes are similar. This was supported by the null results obtained with the cross-classification of category patterns between high and low competition. The lack of significant clusters in the diagonal of the matrix, with only small scattered clusters on the time generalization matrix, suggests that the anticipated category coding in each of the contexts was not alike. Overall, this indicates that although the fidelity of the anticipated content may be the same in high and low competition contexts, the underlying patterns are not shared across conditions, implying a partially different format of preparation depending on the context. Future studies are needed to further confirm this possible explanation. On this respect, task demands could influence how the anticipated information is represented to adapt to the context of the incoming target (Peñalver et al., 2023).

Analysis of the overlap between activity patterns from the perceptual localizer and preparation interval of the main task allowed to examine whether the nature of the category-specific anticipation was similar to perceptually driven patterns (Kaplan et al., 2015; Palenciano et al., 2023). Results showed similarities between perceptual and preparatory category patterns in high competition. Few small clusters were found in low competition. However, there was no statistical evidence of differences between both conditions, which could be due to lack of power to detect small differences in accuracy. Regarding high competition, the overlap between preparatory and visual templates could be associated with the activation of perceptual regions during preparation, as a perceptual reinstatement (Kerrén et al., 2018; Muckli et al., 2015; Smith & Muckli, 2010; Vetter et al., 2014). The partial similarities between preparatory templates and perceptual ones, in the anticipation of high stimuli competition, constitute a relevant finding that contributes to a better understanding of the internal preparatory templates.

Turning to oscillatory activity, how anticipation of competition affects this activity had not been explored in detail in the past. Theta power has been repeatedly related with effort or cognitive control (Cavanagh & Frank, 2014; Cohen & Donner, 2013). Previous studies found that preparing for a difficult task in which stimuli competition is high (Van Driel et al., 2015) or that requires goal updating (Cooper et al., 2017) also induces an anticipatory increase in theta power. Relatedly, and in accordance with our hypothesis, we found that preparing for high competition generated increased theta power in all the preparation interval. Interestingly, this large cluster started right at the beginning of the cue presentation. While this could reflect some extent of smearing of the signal induced by the time-frequency decomposition, this would be unlikely given the high temporal precision for estimating low frequencies such as theta, as the number of cycles assigned to these bands is quite low. Instead, it could be driven by the blocked design employed, which facilitated maintaining the different competition control settings over several trials (Dosenbach et al., 2008; Palenciano et al., 2019b). This temporal profile contrasts with the category-specific anticipation patterns, that are decodable only at the end of the window.

Importantly, our results show that these two different preparatory mechanisms are not correlated. This finding, together with the previously described results, suggest that they reflect different proactive processes that contribute distinctively based on the task requirements. Theta power could implement more general control signals, associated with the general level of competition, whereas category preactivations are specific to the content anticipated (Weber et al., 2024).

Regarding the relationship between anticipatory neural patterns and behavioral performance, some studies suggest that the better these indices, the better task performance (González-García et al., 2017; Manelis & Reder, 2015; Peelen & Kastner, 2011; Soon et al., 2013; Stokes et al., 2009). Our results, however, show inconclusive evidence on this respect. Neither preparatory patterns coding the competition level nor the selected category correlated with behavioral measurements. This was also the case for the correlations with the behavioral congruency effect. This may suggest that the fidelity with which the brain preactivates specific categorical templates of the target does not have a direct influence on the efficiency of behavior, which is in contrast with other studies (González-García et al., 2017; Manelis & Reder, 2015; Peelen & Kastner, 2011). However, the current paradigm was not tailored to entail a wide range of decoding variability in the results, which could hinder the detection of associations between subtle neural preactivations and behavioral measures. Further research is necessary to further explore these associations.

Results show that the neural mechanisms reflected by theta power, in contrast, play an important role for behavioral performance. The effect of competition on RT is partially caused by power in the theta band, indicating that high competition is related to higher theta power which in turn partially explains responses. This finding is in line with previous studies on cognitive control (Cohen & Donner, 2013; Formica et al., 2022). A significant distinction in analyzing the relationship between theta and behavior, compared to the decoding-behavior relation, lies in the utilization of trial-by-trial data in the theta analysis. Obtaining a single value per trial, especially when employing cross-classification to study category-specific patterns, is challenging in the decoding analysis. Other studies have extracted d-values to be more precise (Kerrén et al., 2018; Linde-Domingo et al., 2019; Ritchie et al., 2015), however as we performed cross-classification across cue shapes, this was not feasible. Further tailored studies could address optimal procedures for extracting trial-wise a d-values for the anticipatory neural patterns without visual confounds.

The present study has limitations that restrict the reach of the findings and may catalyze further investigations. First, our scope was the temporal domain, therefore we focused our analyses on the temporal profile of preparatory activity. Further studies may complement our findings with spatially resolved techniques, increasing the anatomical specificity of the different preparatory mechanisms according to competition levels. Moreover, additional studies could use more diverse stimuli over faces and names, and investigate the role of differential difficulty across stimulus categories, which may interact with the competition effect (e.g. Zhang et al., 2013).

Although the current study is not optimized to address this issue, our AUC metric avoids any bias towards a particular stimulus type. Related to this, it may be of interest including multisensory stimuli such as visual and auditory combinations, to replicate and extend the findings to other sensory domains. Furthermore, naturalistic contexts that include different levels of competition could be key to transfer the results to the real environment (Graumann et al., 2022, 2023). Lastly, although our study focuses on preparatory activity, it would be interesting to explore the relationship between attentional templates and target processing. This was not possible on the current dataset because the target-distractor display was substantially different across competition conditions. Future studies could address this issue by reducing the visual differences between conditions. This kind of experiment would enable the examination of the roles played by preparatory attentional templates and anticipatory theta power during actual target processing.

### 4.1. Conclusion

Overall, our results provide insights into how preparation differs depending on the difficulty of the competition that is anticipated. The levels of competition exert a proactive influence on multivariate neural patterns and theta activity. Moreover, the neural mechanisms underlying theta oscillatory activity impact the efficiency of behavior. Integrating these findings into theoretical models of selective attention is crucial for a comprehensive understanding of top-down processes across contexts.

## Author statement

**Blanca Aguado-López**: Data curation, Formal analysis, Writing -original draft. **Ana F. Palenciano**: Formal analysis, Writing -review & editing, Supervision. **José M.G. Peñalver**: Conceptualization, Methodology, Software programming, Data curation, Formal analysis. **Paloma Díaz-Gutiérrez**: Conceptualization, Methodology, Software programming. **David López-García**: Formal analysis. **Chiara Avancini**: Formal analysis. **Luis F. Ciria**: Formal analysis. **María Ruz**: Conceptualization, Methodology, Resources, Writing -review & editing, Supervision, Project administration, Funding acquisition.

## Open practices section

The original data, task, and analysis scripts are openly available. Original code for the EEG preprocessing has been posted on our team Github (https://github.com/Human-Neuroscience/eeg-preprocessing). Raw EEG data, organized following the BIDS format (Pernet et al., 2019) will be deposited in the OpenNeuro public server once the paper is published. The analysis code, results and task code are shared in an Open Science Framework repository that will be public once the paper is published.

## Acknowledgements and fundings

This research was supported by grants PID2019-111187GB-100, PID2022-138940NB, funded by MICIU/AEI/ 10.13039/501100011033 and FEDER, UE, awarded to MR. BAL is supported by a scholarship from the Spanish Ministry of Universities (FPU20/01980). AFP was supported by Grant PAIDI21_00207 of the Andalusian Autonomic Government. JMGP was supported by a scholarship from the Spanish Ministry of Universities (FPU18/01853). CA was supported by a postdoctoral fellowship by the Spanish Ministry for Science and Innovation (FJC2020-046310-I). We are grateful to Christopher I. Williamson for his assistance in data collection.

## Supplementary material

**Fig. S1.**
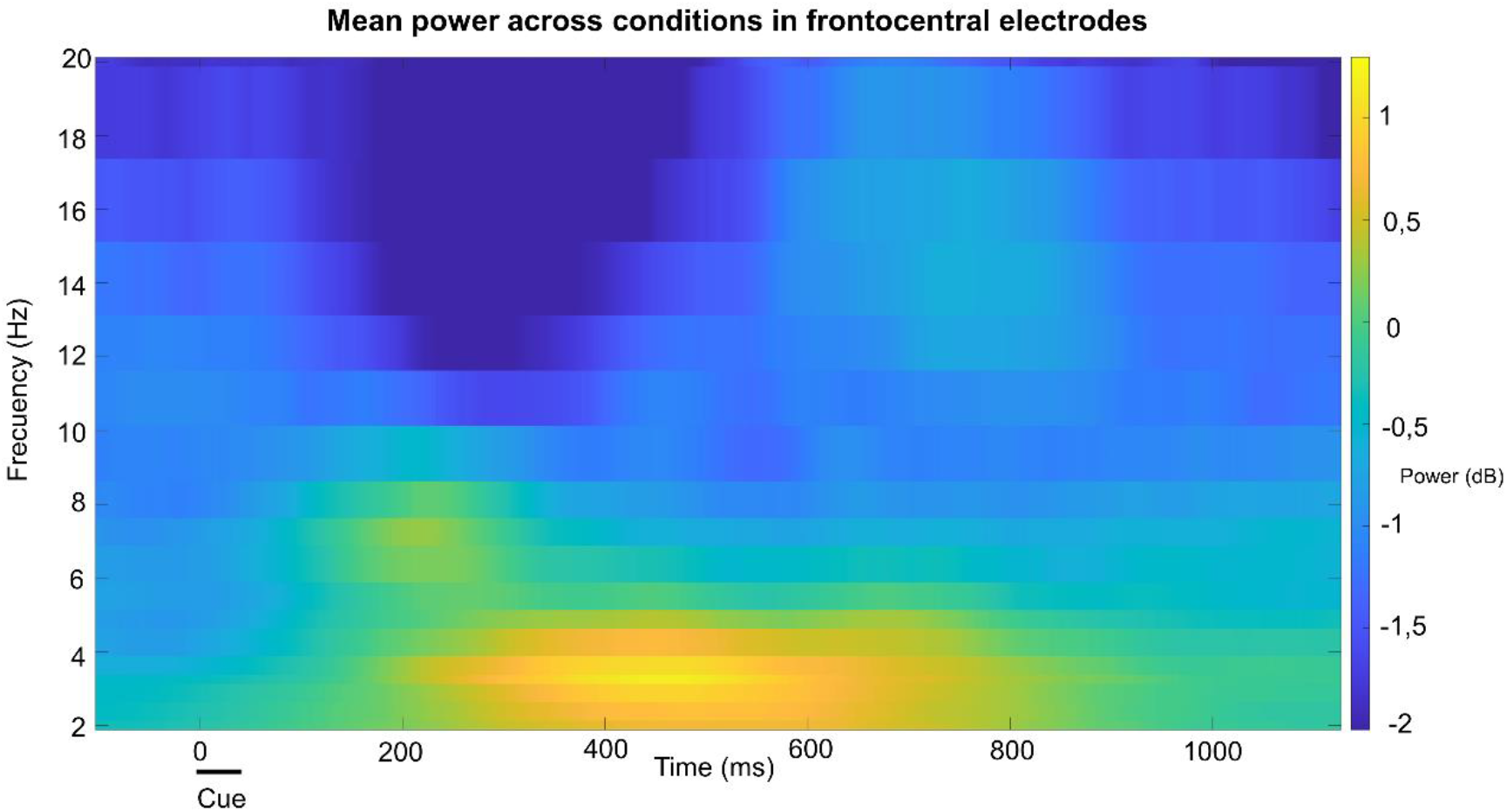
Results of the time-frequency analysis averaged across conditions and participants during the cue-locked window, from -100 ms to 1125 ms.

**Fig. S2.**
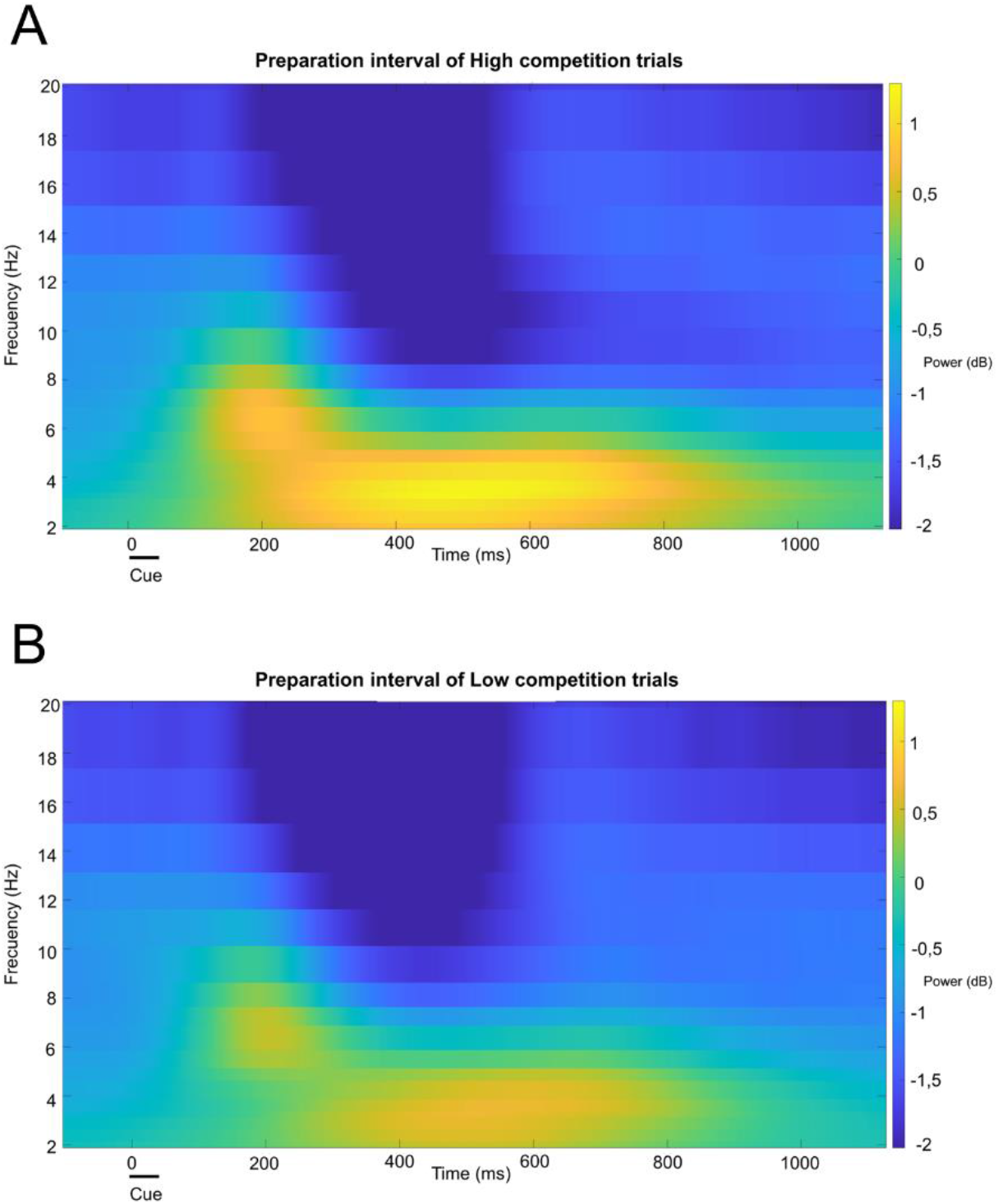
Results of the time-frequency analysis in the preparation epoch of High (A) and Low (B) competition trials from -100 ms to 1125 ms, averaged across participants.

## Supplementary results: LMM equations and results

In our first model we tested if, controlling for congruency, there was an effect of task on RTs, putting a random slope for competition and a random intercept for each participant (RT ∼ competition + congruency + (competition | subject)). In agreement with previous analysis, Low competition blocks were responded faster than High competition ones (t_35.12_ = 4.769, β = 0.021, CI 95% = [0.012, 0.030], *p* < 0.001). Congruency was also significant (t_30807_ = -15.189, β = - 0.018, CI 95% = [-0.020, -0.015], *p* < 0.001). Second, we tested if there was an effect of competition on theta, by fitting an LMM with the same structure but predicting the trial-wise theta values (theta ∼ competition + congruency + (competition | subject)). The effect of competition on theta was also significant, so that High competition induced higher theta power (t_34.7_ = 3.793, β = 0.19, CI 95% = [0.092, 0.289], *p* < 0.001). Third, we tested the fixed effects of theta and competition on RTs, also controlling congruency with the same random structure (RT ∼ theta + competition + congruency + (competition | subject)). The effect of theta was significant (t_30824.2_ = -2.86, β = -0.001, CI 95% = [-0.001, -0.0002], *p* < 0.01), suggesting that larger theta values were associated with faster responses. There was also a main effect of competition (t_35.1_ = 4.807, β = 0.021, CI 95% = [0.013, 0.030], *p* < 0.001) and an effect of congruency (t_30806_ = -15.187, β = -0.018, CI 95% = [-0.02, -0.015], *p* < 0.001).

